# The timescale and functional form of context-dependence during human value-learning

**DOI:** 10.1101/2024.02.01.578398

**Authors:** Maryam Tohidi-Moghaddam, Konstantinos Tsetsos

## Abstract

The way humans and other animals represent the values of alternatives can be distorted by inferior alternatives that are immediately available for choice (immediate context); or that were encountered in previous episodes (temporal context). Yet, the extent to which the immediate and temporal context (co-) shape such context-dependent value coding remains unclear. Here, we asked human participants (N = 30) to learn the values associated with three alternatives and to explicitly report their learned values before making binary and ternary choices among the alternatives. We show that context-dependent value coding is evident in the pre-choice value estimates and manifests equally in binary and ternary choices. Accordingly, we conclude that value representations are modulated by the temporal (and not the immediate) context. Interestingly, the functional form of context-dependence we report runs against extant value normalization theories and is best captured by a new memory-based valuation mechanism we propose.

## Introduction

Does the way you regard two highly desirable products (3^rd^ generation vs. 2^nd^ generation wireless earbuds) change when the seller introduces a third less desirable product (wired earbuds)? Normative theories posit that the relative preference between two alternatives should be independent of the value and the number of other alternatives that are present in the choice set (Luce, 1959, 1977). However, contrary to this premise, empirical findings have shown that the choice between a high-value (HV) and a lower value (LV) alternative can be affected by adding a third distractor alternative (DV) whose value is lower than LV. This phenomenon is known as the distractor effect (Chau et al., 2020; Gluth et al., 2020; Itthipuripat et al., 2015; Louie et al., 2013). The distractor alternative can impact choice quality either in a positive or in a negative manner. In the *positive distractor* effect the tendency to choose HV over LV increases as the value of the DV increases (Chang et al., 2019; Chau et al., 2014, 2020). By contrast, in the *negative distracto*r effect a higher DV alternative reduces the tendency to choose HV over LV (Itthipuripat et al., 2015; Kohl et al., 2023; Louie et al., 2013; Webb et al., 2020).

Distractor effects challenge the independence from irrelevant alternatives principle (IIA) (Luce, 1959, 1977) and pose constraints on the underlying mechanisms that mediate valuation and choice. Random utility models, that apply noise on veridical (unaffected by the context) representations of choice alternatives, predict either a null distractor effect (i.e., logit models) or a feeble positive distractor effect (i.e., probit models, Figure 1B, left) (Berkowitsch et al., 2014; Greene, 2000; Paetz & Steiner, 2018). However, more sizable distractor effects cannot be captured by random utility models. Instead, stronger distractor effects necessitate mechanisms that distort the values of the alternatives based on quantities that relate to the values encountered in a given context (Holper et al., 2017; Khaw et al., 2017; Louie & De Martino, 2014; Rangel & Clithero, 2012; Rigoli, 2019; Webb et al., 2020).

**Figure 1.**
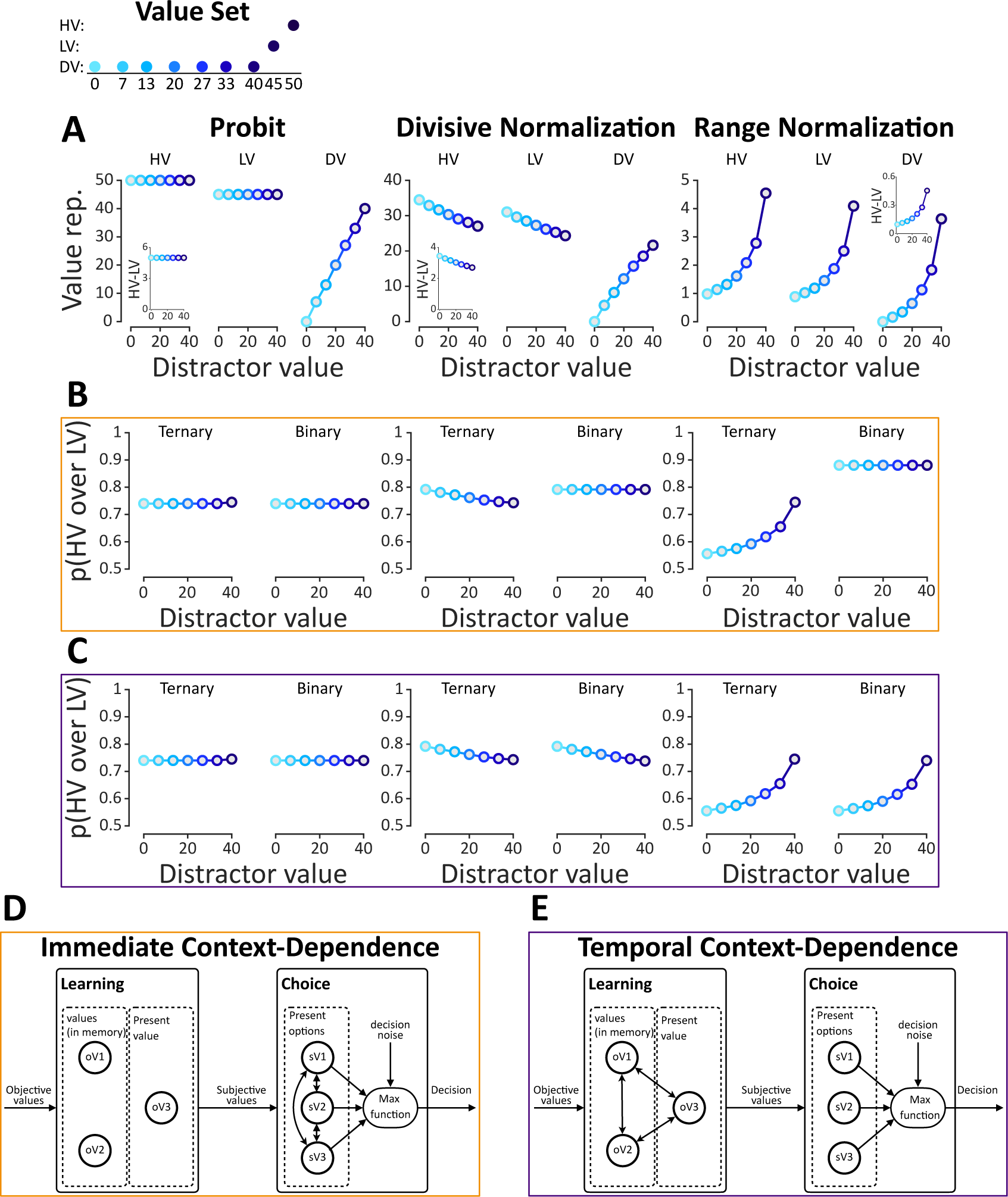
Computational models and predictions. **(A)** Value representations for each alternative by probit (left), divisive normalization (middle) and range normalization (right) models. The high-value (HV) and low-value (LV) alternatives were kept fixed and the distractor value (DV) changed. As the distractor value increases the probit model maintains a constant value for two high-value alternatives (HV and LV), a divisive normalization model predicts a decrease of HV and LV values, and a range normalization predicts an increase of HV and LV values. The embedded plots depict the difference between the value representation of HV and LV as a function of the distractor value. **(B)** Simulated relative choice of HV over LV in the “immediate-context” framework and **(C)** “temporal-context” frameworks. In each subplot, *P(HV over LV)* is calculated separately for ternary (HV, LV, DV) and binary choice trials (HV, LV). **(D)** Visual depiction of the immediate and **(E)** temporal context-dependence frameworks. The two-sided arrows indicate computations leading to context-dependent distortions between alternatives. One-sided arrows indicate the transfer of information to the corresponding node. (for further details, see Supplemental Material)

In particular, in the range normalization model, each alternative value is divided by the range of all values (i.e., maximum *minus* minimum) in the current context (Kontek & Lewandowski, 2018; Palminteri et al., 2015; Soltani et al., 2012). Therefore, when the context features a higher distractor value, it leads to a narrower range or normalization term. This gives rise to a higher distorted value (i.e., subjective value) for both of the two high-value alternatives (Figure 1A, right), and a greater difference between them (Figure 1A, right, embedded plot), effectively leading to a positive distractor effect (Figure 1B, right). By contrast, in the divisive normalization model, each value is normalized by the sum of all values in the context (Louie et al., 2013). Thus, a higher distractor value results in a larger normalization term, dwarfing both the distorted values for the two high-value alternatives (Figure 1A, middle) and the value difference between them (Figure 1A, middle, embedded plot). Consequently, this type of distortion gives rise to a negative distractor effect (Figure 1B, middle).

Despite the recent proliferation of experimental studies and the development of normalization models focusing on context-dependence, definitive conclusions on the underlying mechanisms remain elusive (Cao & Tsetsos, 2022; Chau et al., 2020; Gluth et al., 2020; Hayes & Wedell, 2023; Juechems et al., 2022; Louie et al., 2013). First, there are controversies around the exact direction and robustness of distractor effects. Specifically, both positive (Chau et al., 2014, 2020; Gluth et al., 2018) and negative (Itthipuripat et al., 2015; Louie et al., 2013) distractor effects have been reported using different experimental paradigms ranging from reward-learning in primates to risky and preferential choice paradigms in humans. Some of these effects failed to replicate (Gluth et al., 2018, 2020) or changed direction after re-analysis (Cao & Tsetsos, 2022; Chau et al., 2020). Second, previous studies have not systematically examined the timescale in which context-based distortions operate. Value representations could be distorted on the basis of the values of alternatives that were previously encountered (temporal context, Figure 1E); and also, on the basis of the values of the alternatives that are immediately available on a given choice trial (immediate context, Figure 1D) (Frydman & Jin, n.d.; Louie & De Martino, 2014; Madan et al., 2021). It is worth mentioning that some context effects arise in one-shot decisions embedded in non-specific temporal contexts (Chau et al., 2014; Gluth et al., 2018, 2020; Louie et al., 2013). In these cases, the immediate context account is the only viable one. However, in tasks that involve learning the values of alternatives within specific temporal contexts (i.e., with each context being characterized by a different distractor value) and subsequently making choices based on these learned values (i.e., subjective values), the distinction between temporal and immediate context is blurry (Bavard & Palminteri, 2023; Hayes & Wedell, 2023; Louie et al., 2013; Palminteri & Lebreton, 2021). The key experiments that could differentiate the influence of the immediate context from that of the temporal context in value-learning tasks are yet to be conducted.

Driven by the above, we here probed distractor effects within a single experimental paradigm that does not tap upon complex cognitive processes evoking alternative interpretations for distractor effects (Chau et al., 2014, 2020). In particular, we used a simple value-learning task previously shown to induce a negative distractor effect in primates (Louie et al., 2013). Like in previous studies, to assess the magnitude of the distractor effect, we varied the value of the distractor alternative (DV) across contexts (experimental mini-blocks). Our design differed from previous studies (Cao & Tsetsos, 2023; Itthipuripat et al., 2015) in the following aspects. First, in our task, the reward values associated with each alternative were independently learned before the choice phase. Second, we interleaved a value estimation task immediately after the learning phase so that we could directly gauge people’s subjective values before the onset of the choice phase. That way we could ask whether value distortions had already occurred at the end of the learning epoch, which would support the temporal context hypothesis (Figure 1C and 1E). Finally, in the choice phase we interleaved binary and ternary choice trials. Comparing the relative choice of HV over LV in these two trial types, we could assess whether the immediate availability of the distractor alternative affects choice (Figure 1B and 1D, immediate context hypothesis).

To outline our results, our data support that value distortions operate on a longer timescale (i.e., the temporal context) while the immediate context had no effect. Specifically, in line with the temporal context hypothesis, we found that subjective value estimates (elicited in the value estimation phase, interchangeably referred to as estimated values) are reduced under high distractor values, already after the learning phase. This reduction applies equally to the two high-value alternatives (HV and LV), a finding that falsifies both divisive and range normalization models. In the choice phase, we obtained a trend towards a positive distractor effect (at first glance consistent with range-normalization) which did not vary between binary and ternary trials. The lack of difference in the distractor effect between binary and ternary trials suggests that distortions in subjective value are not influenced by the immediate context but rather take shape over a longer timescale. Although, the subjective value estimates participants provided after learning predicted their subsequent choices, the functional form of the distortion was different in the two phases (i.e., a global reduction in value estimates vs. a trend toward a positive distractor effect, respectively). Inspired by the decision-by-sampling framework (Noguchi & Stewart, 2018; Stewart et al., 2006), we outline a stochastic rank-based mechanism that potentially reconciles this discrepancy. Taken together, our results shed new light on the timescale and computational mechanisms that underlie context-dependent multialternative choices.

## Methods

### Participants

We recruited 30 participants (22 female, *M* = 25.3 years, *SD* = 3.41) from the internal participant pool in the Institute of Neurophysiology and Pathophysiology of the University Medical Center Hamburg-Eppendorf. Before starting the experiment, all participants gave written informed consent. The study was approved by the local ethics committee of the Hamburg Medical Association and conducted in accordance with the Declaration of Helsinki. All participants had normal or corrected-to-normal vision were right-handed, with the exception of one participant who was left-handed. The experimental session lasted between 3 and 3.5 hrs. Participants received 30-35 EUR for their participation (hourly net rate = 10 EUR). In addition, we included a 10 EUR completion bonus and a maximum of 15 EUR task performance bonus to further incentivize participants to accurately perform the task (for further details, see Supplemental Material).

### Procedure

Participants were instructed to complete 48 mini-blocks of a value learning task (Figure 2A). Their task was to collect valuable-colored coins to maximize their overall obtained reward. Each mini-block started with a learning phase where participants learned the association between colored coins and reward-values (Figure 2B). Later, they faced binary and ternary choices in the choice phase, aiming to choose the alternative (colored coins) with the highest reward (Figure 2D). Following the learning and choice phases we added two estimation phases, in which participants reported their subjective value estimates for each colored coin (Figure 2C). In each mini-block, the value of all three alternatives (i.e., the context) was kept fixed. We randomly assigned the 3 alternatives (HV, LV, DV, specific to each mini-block) to three colored circles (red, blue, and yellow), mapping them onto three imaginary coins of different values. In total, we created four unique contexts by deploying two levels of distractor DV = {18, 40} allowing us to assess the distractor effect, and two magnitude levels to insert variability into the task HV= {55, 50}, LV = HV – 5 (The magnitude level (HV equal to 55 or 50) did not yield any significant effects in any of the key analyses. Therefore, for the sake of simplicity, we collapsed the data across the two magnitude levels.). Combining these four contexts with a full color-reward permutation, led to 24 unique possible mini-blocks (six color-reward permutations times four unique value contexts). These mini-blocks were repeated twice, in a total of 48 mini-blocks. In one-half of the mini-blocks, we did not provide any reward feedback in the choice phase (“No-Feedback” mini-blocks) while in the other half we showed the reward associated with each choice (“Feedback” mini-blocks). We verbally instructed participants that “Feedback” and “No-Feedback” mini-blocks were randomly interleaved and that there was no way to know which mini-block they were performing prior to the choice phase. This ensured that the participants paid attention to the learning phase equally in “Feedback” and “No-Feedback” mini-blocks, enabling us to attribute any behavioral disparities across the two conditions to the choice phase (for further details, see Supplemental Material).

**Figure 2.**
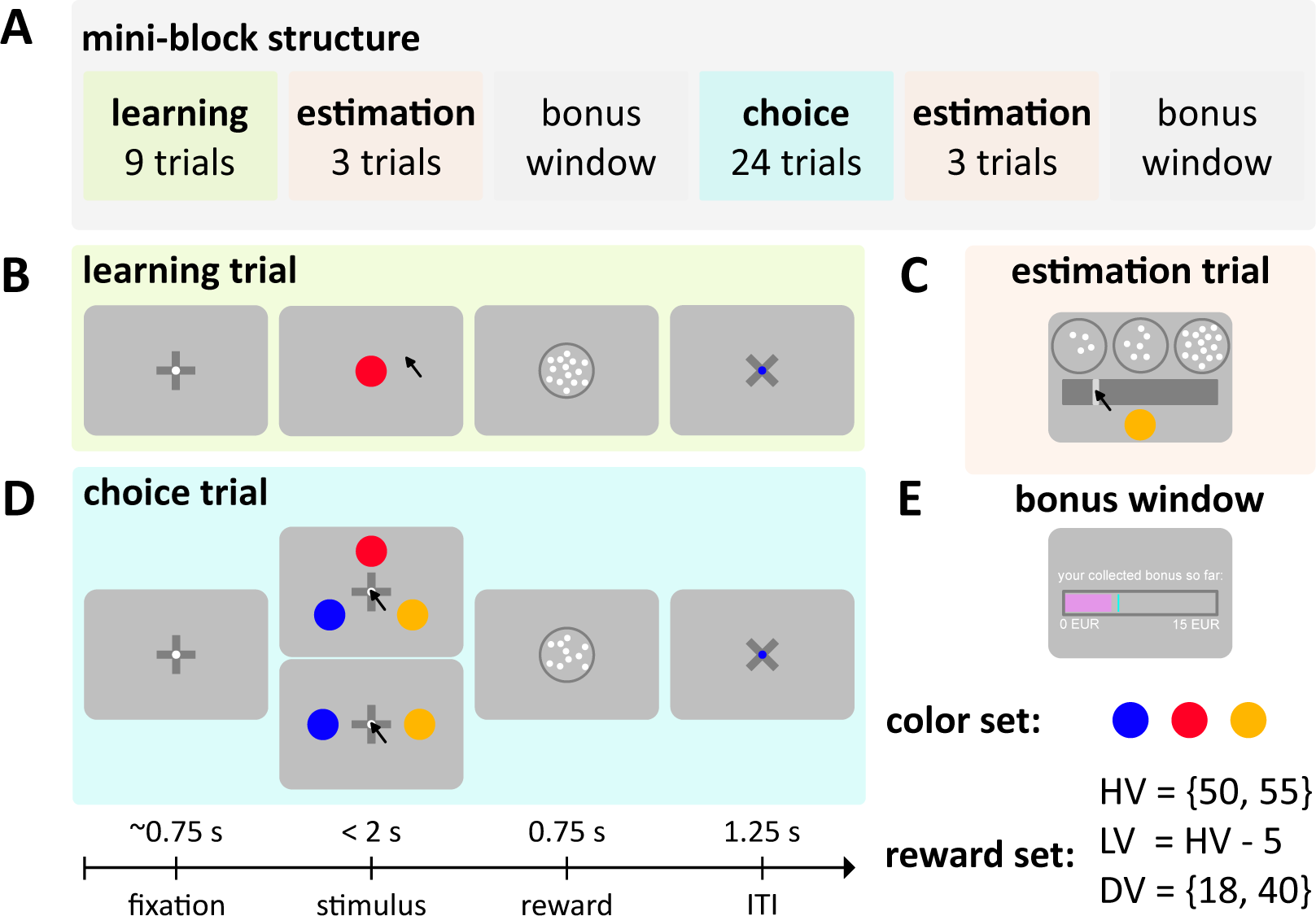
Human value learning task. **(A)** Mini-block timeline (from left to right), each participant completed 48 mini-blocks of color-reward association task. They were instructed to maximize their total obtained reward by first learning the color-reward association during the learning phase of each mini-block and then choosing the alternative with the highest reward in the subsequent choice phase. Each mini-block started with the learning phase where participants sequentially observed the values associated to 3 alternatives (imaginary colored coins, look at the color set) in a given context. Each context included a unique HV, LV, and DV alternative (c.f. reward set). Following the learning phase, each mini-block continued with the first estimation phase where participants reported their subjective value estimate of each alternative. After the estimation phase, the bonus window shortly appeared before the choice phase began. In the choice phase participants had to choose the alternative that yielded the largest reward. The choice phase was followed by the second estimation phase and each mini-block ended with the presentation of the final bonus window. **(B)** The learning phase consisted of 9 learning trials (3 per each alternative). Each trial began with the fixation cue for a random duration, and after that one of the alternatives appeared on the screen. Participants were asked to click on the alternative within 2 seconds to get the associated reward (i.e. represented by the number of a set of moving dots). **(C)** The Estimation phase involved 3 trials (1 per each alternative). On each trial, an alternative was presented below a slider bar and participants were asked to move the slider to indicate the number of dots (i.e., reward value) associated with that alternative. **(D)** The choice phase included 24 trials (12 ternary and 12 binary). Like learning trials, each choice trial initiated with the fixation cue for a random delay, and after that, 2 or 3 alternatives presented on the screen. Participants were asked to click on their preferred alternative within 2 seconds to see the collected reward. In “No-Feedback” mini-blocks, the choice was immediately followed by the inter-trial-interval (ITI) window, and the collected reward was not presented. **(E)** The bonus window, presented after each estimation phase, revealed the participant’s collected monetary bonus in a pink bar and the possible maximum collected bonus with a cyan line so far. (for further details, see Supplemental Material)

### Analyses

Across different analyses, the statistical effects were examined using one-sample *t*-tests, *F*-test using ANOVA, and Pearson correlation. In the correlation analyses, we first obtained a correlation coefficient for each participant and then compared the correlation coefficients of the group against zero using a two-sided *t*-test on. All analyses were done using MATLAB (The MathWorks Inc., 2022) and we used JASP (JASP Team, 2023) for Bayesian *t*-tests.

To quantify the distractor effect, we compared the relative choice in the choice phase between the two contexts with the high-value and low-value distractors. The relative choice is quantified as:

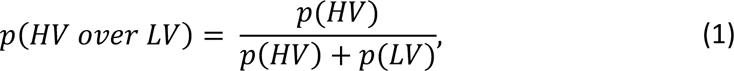

where *p(HV)* and *p(LV)* indicate the choice probability of the high-value and low-value alternatives. Then, the distractor effect is quantified as:

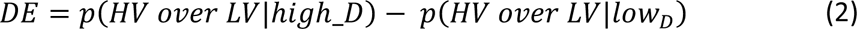

Thus, *DE > 0*, *DE < 0*, and, *DE = 0*, indicate a positive, a negative, and a null distractor effect respectively. We only analyzed trials in which participants responded within the 2-second deadline in the learning and choice phases and the 10-second deadline in the estimation phases (for further details, see Supplemental Material).

## Results

### Basic performance

During the learning phase (Figure 2B), participants were sequentially exposed to the value associated with each alternative after clicking on a centrally presented colored coin within a maximum of two seconds. After clicking, a cloud of moving dots briefly appeared, with the number of dots representing the reward magnitude (i.e., number of points) associated with the coin (see Supplemental Material). Participants completed 99.91% (95% CI = [99.84%, 99.97%]) of all learning trials with an average reaction time of 623.66 milliseconds (ms) (95% CI = [593.56 (ms), 653.76 (ms)]), indicating that they properly engaged with the learning phase. We did not see any differences between the reaction times across learning trials that involved the HV, LV, and DV alternatives across different contexts (2-way ANOVA on *log*(RT), alternatives: *F*(2,179) = .004, *p =* .99, contexts: *F*(1,179) = 0.15, *p =* .69, interaction: *F*(2,179) = 0.02, *p =* .98).

In the choice phase (Figure 2D), participants made 12 ternary and 12 binary choices (four trials per each possible binary pair) by clicking on their preferred alternative within a maximum of two seconds. We included all three possible binary pairs (HV vs. LV, HV vs. DV, LV vs. DV) to keep the presentation rate of all alternatives equal and to avoid introducing any implicit bias toward the two high-value alternatives. Participants completed 99.73% (95% CI = [99.62%, 99.84%]) of all choice trials with an average reaction time of 729.89 (ms) (95% CI = [695.14 (ms), 764.65 (ms)]). Participants chose the best alternative with 81.63% (95% CI = [78.49%, 84.77%]) accuracy in ternary (chance level 33.3%) and 90.27% (95% CI = [88.29%, 92.25%]) in binary trials (chance level 50%; one-sided *t*-test against chance level on ternary: *t*(29) = 30.17, *p* < .001, and on binary: *t*(29) = 39.86, *p* < .001). The average reaction time in binary trials was significantly higher than in ternary trials and did not change across different contexts (2-way ANOVA on *log*(RT), choice type: *F*(2,119) = 11.40, *p* < .001, contexts: *F*(1,179) = .20, *p =* .65, interaction: *F*(1,179) = .03, *p* = .87, see Figure S1 in Supplemental Material). This seemingly counterintuitive finding might be attributed to the greater uncertainty linked with the identities of the alternatives in binary trials.

Moreover, in half of the mini-blocks, we withheld reward feedback in the choice phase to intercept further learning from feedback after the value estimation phase. Withholding feedback could help us better quantify immediate context effects. We found that receiving feedback did not significantly improve choice accuracy (choice accuracy pooled across both ternary and binary: in “Feedback”, 89.00% (95% CI = [86.25%, 91.76%]) and in “No-Feedback”, 87.22% (95% CI = [85.02%, 86.42%]) mini-blocks; two-sided *t*-test: *t*(29) = 1.54, *p* = .13). The average reaction time was not influenced by receiving feedback (reaction time pooled across both ternary and binary: in “Feedback”, 730.46 (ms) (95% CI = [696.85 (ms), 764.07 (ms)]) and in “No-Feedback”, 729.36 (ms) (95% CI = [691.98 (ms), 766.75 (ms)]) mini-blocks; two-sided *t*-test on *log*(RT): *t*(29) = .39, *p* = .70). Overall, the choice rate (i.e., the number of times one alternative is chosen *divided by* the number of times that alternative is presented) of each value was consistent with value-guided decision-making (choice rate of (HV) = 84.83%, (95% CI = [82.17%, 87.49%]); (LV) = 31.80%, (95% CI = [29.94%, 33.65%]); and (DV) = 3.32%, (95% CI = [2.08%, 4.56%])), pooled across both “Feedback” and “No-Feedback” mini-blocks).

In the estimation phase (Figure 2C), participants were asked to report their subjectively learned value of each alternative by moving a slider on a bar within 10 seconds. Participants completed 99.70% (95% CI = [99.51%, 99.89%]) of all estimation trials across both phases with an average reaction time of 3.24 seconds (sec) (95% CI = [2.95 (sec), 3.52 (sec)]). In both estimation phases, the average estimated values (i.e., subjective values) correlated strongly with the actual values (Pearson correlation, First estimation: *r* = .984 (95% CI = [.979, .990]), two-sided *t*-test on correlation coefficients*: t*(29) = 334.17, *p* < .001, Second estimation: *r* = .98 (95% CI = [.97, .99]), two-sided *t*-test on correlation coefficients*: t*(29) = 251.69, *p* < .001). All analyses above indicate that participants engaged well with the estimation and choice tasks.

### Context-dependent distortions during learning

Having ensured that participants complied with the task instructions, we next asked how value estimates (i.e., subjective values) varied as a function of the distractor value. Here, we analyzed the estimation data pooled across both “Feedback” and “No-Feedback” mini-blocks (as this manipulation was announced to the participants before they started the choice phase, see Method). We only focused on the results from the first estimation as these were not influenced by any behavior or feedback at the choice phase. Figure 3A shows the average value estimates for each alternative across contexts. The data reveal that the estimated values of the two high-value alternatives significantly decreased in the high-distractor context relative to the low-distractor context (two-sided *t*-test in First estimation on HV: *t*(29) = 5.33, *p* < .001, on LV: *t*(29) = 5.38, *p* < .001). This effect indicates that distractor-induced distortions in subjective value already occurred during the learning epoch.

**Figure 3.**
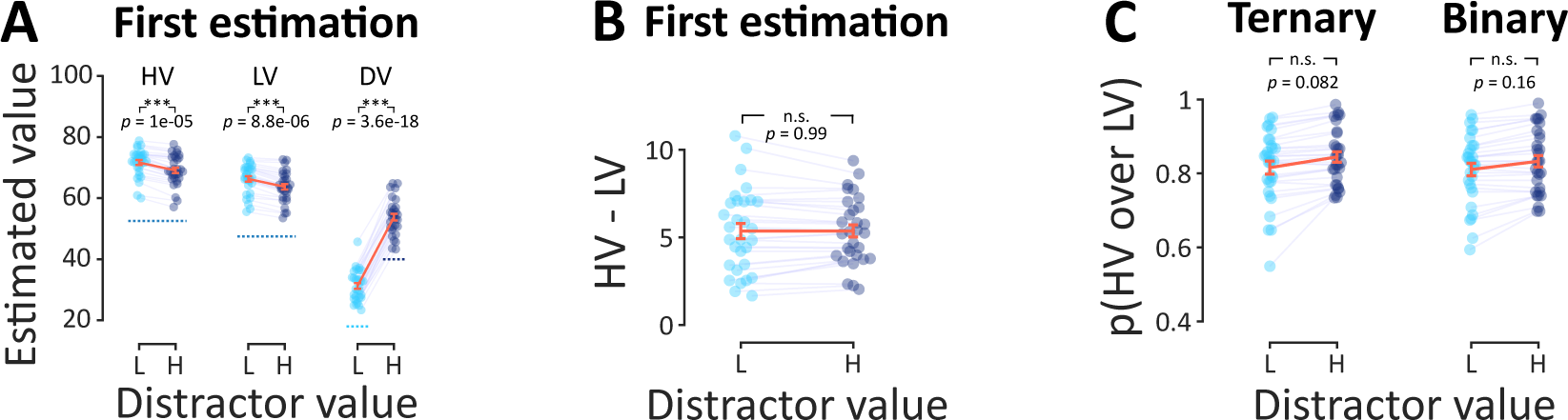
Context dependence in estimated value and choice. **(A)** Mean estimated value of each alternative as a function of the distractor value. Dashed lines indicate the actual values. **(B)** The difference in estimated values between the two high-value alternatives, shown in the two distractor contexts. **(C)** Relative choice of HV over LV in the choice phase as a function of the distractor value in ternary trials and binary (HV, LV) trials. In all panels, data were pooled across “Feedback” and “No-Feedback” mini-blocks, the orange error bars indicate standard errors of the mean across participants, and the colored dots indicate individual participants.

At first glance, this pattern appears consistent with the divisive normalization prediction and runs against the range normalization prediction (Figure 1A). However, divisive normalization also predicts that the value difference between HV and LV decreases as the distractor value increases (embedded plot in Figure 1A, middle). Contrary to this prediction, figure 3B shows that the value difference stays the same in both contexts (two-sided *t*-test on “*HV – LV*” between the contexts with low and high distractors, First estimation: *t*(29) = -.006, *p* = .99). This null finding in the value difference is more consistent with a probit model, which does not predict a global reduction though, and inconsistent with both normalization models (embedded plot in Figure 1A, left).

### A trend for a positive distractor effect in both binary and ternary choice trials

We next sought to assess whether the choice data revealed a distractor effect. To probe the distractor effect we compared the relative choice share between the two high-value alternatives (here denoted as *p(HV over LV)*, see Equation 1) in ternary choices across high-distractor vs. low-distractor mini-blocks (Fig 3C, left labelled as “Ternary”). In contrast to the prediction of the divisive normalization and the findings in Louie et al. (Louie et al., 2013), we found a trend for a positive distractor effect (two-sided paired *t*-test on Ternary: *t*(29) = -1.8, *p* = .082) which is consistent with the range normalization model (Figure 1C, “range-normalization” prediction). Here the data is pooled across both “Feedback” and “No-Feedback” mini-blocks, given that there was no difference in the distractor effect between these two mini-block types (two-sided paired *t*-test: *t*(29) = -.44, *p* = .66).

Given the absence of a negative distractor effect and the trend for a positive distractor effect we independently quantified the evidence in favor of either of these two opposing effects relative to a null hypothesis where no distractor effect occurs. We conducted two one-sided *t*-tests within a Bayesian framework with the respective alternative hypotheses. The test featuring the negative distractor effect as the alternative hypothesis provided a Bayes factor (BF_10_) of 0.08 (95% CI = [0.002, 0.24]), indicating strong evidence in favor of the null hypothesis. On the other hand, the test with the positive distractor effect as the alternative hypothesis resulted in a Bayes factor (BF_10_) of 1.39 (95% CI = [0.033, 0.65]), suggesting anecdotal evidence in favor of a positive distractor effect. We additionally fitted the choice data and performed a model-based comparison between probit, divisive normalization, and range normalization, as these models predict a nearly zero, a negative, and a positive distractor effect, respectively. According to a fixed-effect comparison, the range normalization fits yielded the minimum BIC score for 67% of the participants, while the probit model could describe the remaining 33%. We further performed a random-effects model comparison using a cross-validation approach (see Supplemental Material). Here, the range normalization was also the winning model with a posterior model frequency equal to 97% (Pexc > .99).

To assess the timescale of context-dependence introduced earlier (also Figure 1B-E), we turned into comparing the choice probabilities between ternary (HV, LV, DV) and binary (HV, LV) trials. If the distractor effect (Equation 2) present in ternary trials is different from the distractor effect in binary trials (Figure 3C, right labelled as “Binary”), then this will support the hypothesis that the immediate context contributes to context-dependent value distortions. If on the other hand, the binary and ternary distractor metrics are indistinguishable, we can infer that the trend for a positive distractor effect is entirely attributable to the temporal context. In accordance with the latter, the distractor effect in ternary and binary choices was indistinguishable (Figure 3C, two-sided paired *t*-test on the distractor effect: *t*(29) = .98, *p =* .34). Additionally, the binary and ternary distractor effects were strongly correlated across participants (Pearson correlation, *r*_(30)_ = .90, *p* < .001) further corroborating the lack of immediate context value modulation. Thus, the consistent binary and ternary choice patterns indicate that the value of the distractor alternative modulates value representations across a prolonged timescale, extending beyond the level of a single choice trial.

### Common value representations in estimation and choice

The estimation data revealed a strong contextual modulation, with high distractor values leading to an overall underestimation of the two high-value alternatives. Importantly, the distractor value did not affect the HV and LV differently in the estimation phase. This subtractive distractor effect runs against the positive distractor trend observed in the choice phase. Here we attempted to better understand why the context effects differ between the estimation and choice phases.

We started with two hypotheses placed at the opposite ends of a spectrum. First, this discrepancy could be driven by completely distinct processes underpinning value representations in the estimation and choice phases. In that case, we would expect that the value estimates are not predictive of the choices people made (H1). Alternatively, there could be a common subjective value representation underlying both tasks, but because the magnitude of the positive distractor effect is very small, reliably detecting an equivalent effect in the estimation phase is not possible if the latter is influenced by greater levels of noise (H2).

To arbitrate between these two hypotheses, we examined the consistency between value estimates and choices. To do so, we applied an established approach (see Equation S1 and S2 in Supplemental Material (Bavard & Palminteri, 2023)) to calculate pseudo-choices that would have been made had participants relied on the values reported in the estimation phases. We then compared the choice rate between the actual choices and the pseudo-choices. Assessing the choice rate allowed us to combine binary and ternary choices in one quantity. Figure 4A illustrates that pseudo-choices obtained from the value estimates are highly correlated with the actual choices (Pearson correlation, First estimation: *r* = .994 (95% CI = [.991, .996]), two-sided *t*-test on correlation coefficients*: t*(29) = 595.62, *p* < .001, with Second estimation: *r* = .992 (95% CI = [.987, .998]), two-sided *t*-test on correlation coefficients*: t*(29) = 363.69, *p* < .001). Notably, the first estimation displayed a closer match than the second estimation (average “mean squared error” of choice rate across the three alternatives in the First estimation = 19.62, (95% CI = [9.01, 30.23]); and in the Second estimation = 27.05, (95% CI = [7.70, 46.41])). We further correlated the “*HV – LV*” difference from the first estimation phase against the distractor effect (Equation 2) quantified in the choice phase when combining binary and ternary trials (Figure 4B, Pearson correlation between the distractor effect and “*HV – LV*” estimate difference between two contexts in the First estimation: *r*_(30)_ = 0.57, *p* < .01, and in the Second estimation: *r*_(30)_ = 0.65, *p* < .001). This significant correlation, in conjunction with the pseudo-choices analysis above, indicates a non-negligible link between the value estimates and choices.

**Figure 4.**
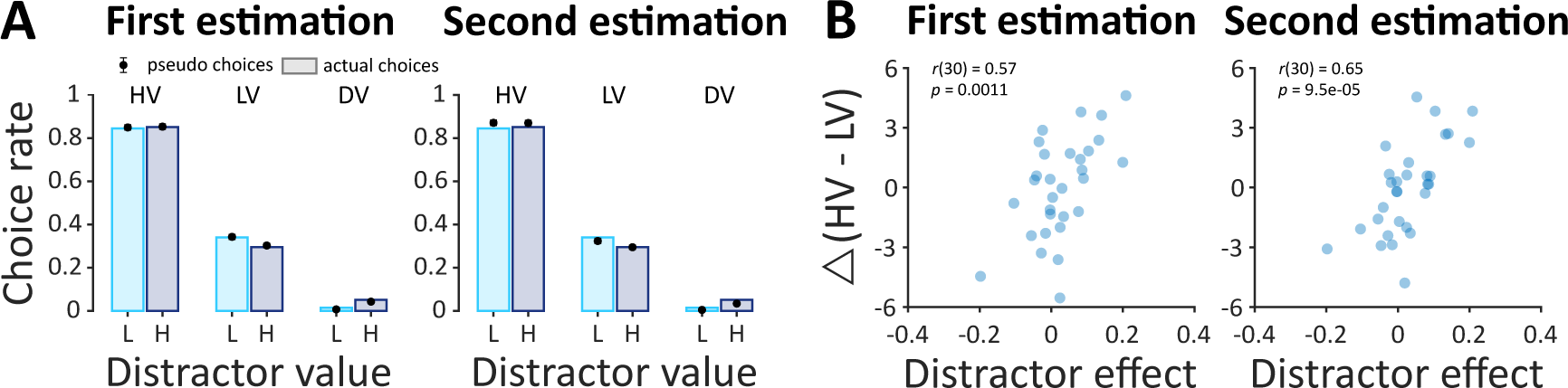
Value representation comparison between the estimation and choice phases. **(A)** Choice rates (bars) obtained in the choice phase and pseudo-choice rates (filled circles) derived from the estimated values (Equation S1 and S2 in Supplemental Material). **(B)** Correlation between the distractor effect (Equation 2) and *“HV – LV”* estimate difference between two contexts. Error bars in A indicate standard errors of the mean across participants and in B the blue dots indicate individual participants.

Given these results, we can therefore rule out H1 according to which value representations in the two tasks are completely unrelated to each other. This motivates us to develop a single model that can jointly explain the context effects observed in the estimation and choice phases. Thus, in the following section we outline a model that can produce a positive distractor effect while reducing the value representations for both high-value alternatives as the value of the distractor increases.

### Context-dependent behavior beyond normalization theories

Overall, none of the two normalization models can adequately capture the observed estimation and choice patterns. Under the assumption that estimation and choice derive from a common underlying value representation (H2 above), a successful model needs to predict a) a strong reduction of HV and LV values under high distractor, b) a (small) positive distractor effect evident in both the “*HV – LV*” estimates and in the choice probabilities.

We propose a rank-based (RB) model (Stewart et al., 2006) that can capture these patterns in both estimation and choice data. This model relies on the decision-by-sampling framework that has been successfully used to predict context effects in various settings, including multi-alternative, multi-attribute choices (Noguchi & Stewart, 2014; Paetz & Steiner, 2018; Stewart et al., 2006). The RB model assumes the value representation of an alternative in a choice trial is constructed via a series of stochastic binary comparisons between memory samples of the target and of previously encountered alternatives (Figure 5A and 5B). Thus, by-definition, this model operates at a long timescale retrieving previously encountered values from the memory context in order to construct the value of a target alternative. The constructed value (i.e., subjective value) of a target alternative is simply the number of times it won in these binary comparisons (Figure 5C). As a result, in a higher distractor context, the constructed values are overall reduced for both HV and LV due to increased competition with DV. The reduction is stronger for LV though, due to its proximity to DV, leading to a higher delta of “*HV – LV*” and subsequently giving rise to a positive distractor effect (Figure 5C).

**Figure 5.**
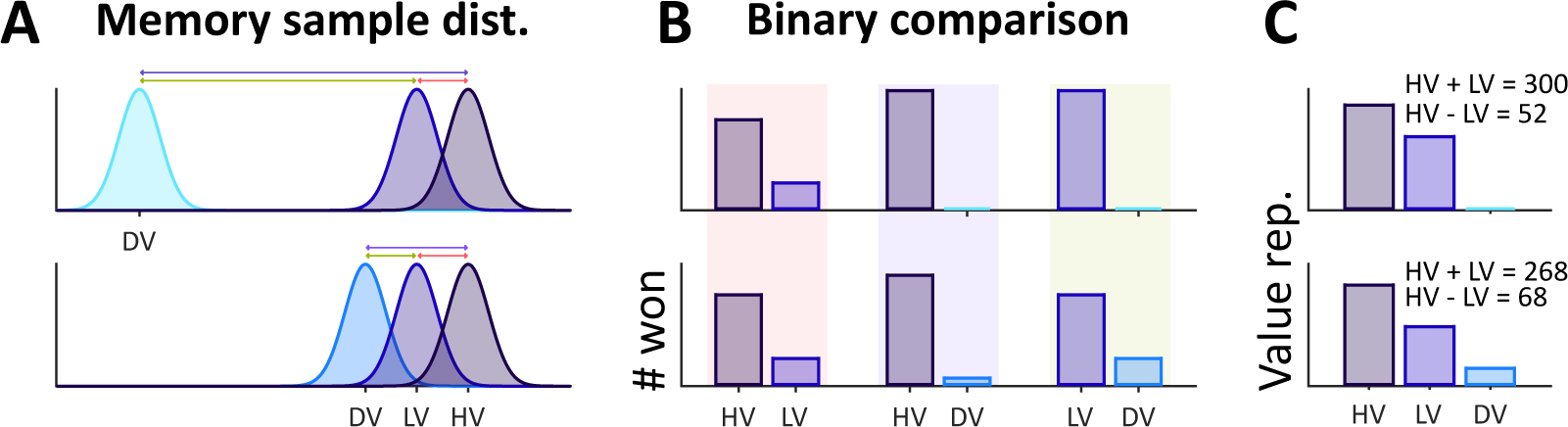
Rank-based (RB) model illustration. (A) The distribution of memory samples of the objective values. The top and bottom row are in the presence of a low and high value distractor respectively. **(B)** The schematic of number of times an alternative won in arbitrary 100 binary comparisons (k parameter in the RB model), here all possible binary comparisons were presented. For example, in the bottom row the HV once compared with LV (indicate by red arrow/shade) and once with DV (indicate by purple arrow/shade). **(C)** The final value representation of each alternative. Here for example the value of HV is the sum of the number of times HV won against LV and DV.

Figure 6A depicts that the RB model provided a closer fit to the choice distractor effect. We show this by simulating choices using the best-fitted parameters of each model, and then pooling the model predictions across both ternary and binary choices given that there was no difference in the distractor effect between ternary and binary choices. Two further model comparisons confirmed that the RB model accounted for the choice data (Figure 6B and 6C) better than the other alternative models. First, in a fixed-effects comparison, the RB model has the smallest difference relative to the best-individualized model (ΔBIC > 10 (Kass & Raftery, 1995)) (RB, Pr, and RN had the smallest BIC for 46.7 %, 26.7%, and 26.7% of the participants and marked as the best-individualized model). Second, in a random-effects model comparison the RB model is better supported (*P*pexc > .99 (Rigoux et al., 2014)) (for further details, see Supplemental Material).

**Figure 6.**
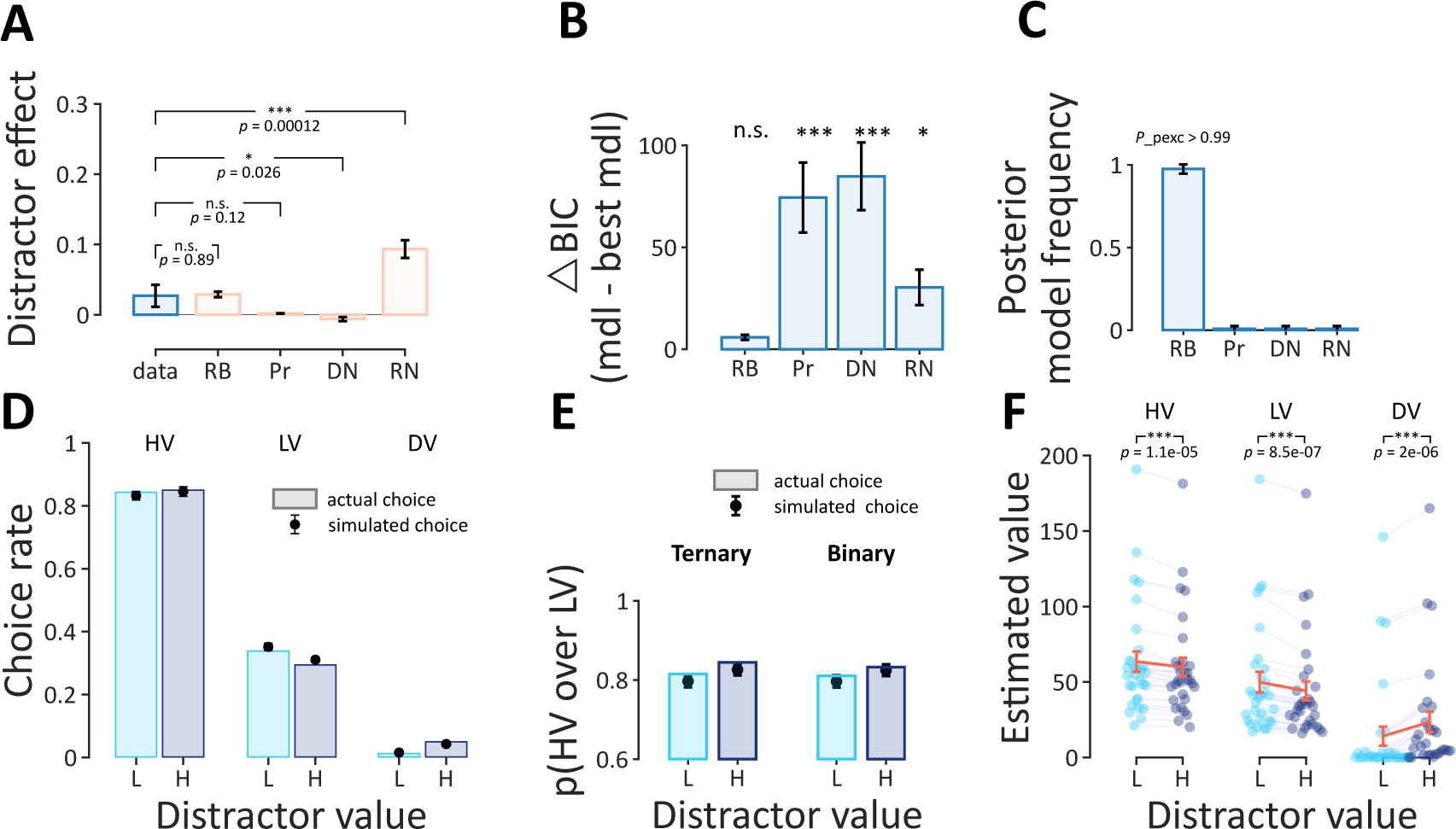
Model comparison. **(A)** Distractor effect comparison between the actual data and simulated data using the best-fitted parameters, the RB model has a closer fit to the actual distractor effect in the choice data. **(B)** Fixed-effect Bayesian model comparison, Black asterisks: significant ΔBIC against 10, one-sided one-sample *t*-tests (Kass & Raftery, 1995). **(C)** Random-effect Bayesian model comparison (Daunizeau et al., 2014). **(D)** The simulated choice rate of the RB model superimposed on the actual choice. **(E)** The simulated relative choice of the RB superimposed on the actual relative choice. **(F)** The estimated values of each alternative, simulated by the best-fitted parameters of the RB model. In all panels, the error bars are standard errors of the mean across participants. In panel F, the colored dots indicate the individual participant.

Figures 6D and 6E show that the RB model can describe the choice rate and relative choice of HV over LV well (see Figure S2 and Table S1 in Supplemental Material for the fits of the other models and the best-fitted parameters of all models, respectively). To demonstrate that the RB model can approximate the estimation data, we extracted the value representations afforded by the model using the best-fitted parameters. Figure 6F demonstrates that the RB model predicts a large decrease in the estimated values of the two high-value alternatives (HV and LV) as the distractor value increased, equivalent to what we observed in the behavioral data (Figure 3A). The model also predicts a larger delta “*HV – LV*” in the estimation data under the high distractor, which was not observed in the data. As H2 in the previous section posits, the lack of “*HV – LV*” effect could occur if we assume a larger amount of noise in the estimation process. Thus, the RB model can reconcile the subtractive distractor effect in estimation and the small positive distractor effect in choice, under a single mechanism.

## Discussion

Previous findings suggest that the value of an inferior (distractor) alternative influences choices between two high-value alternatives (Chau et al., 2020; Louie et al., 2013; Louie & Glimcher, 2012). However, the functional form of the distractor influence remains generally contentious while its timescale in value-learning tasks is underexplored (Frydman & Jin, n.d.; Louie & De Martino, 2014; Madan et al., 2021). Using a value-learning paradigm that was previously shown to induce a negative distractor effect in primates (Louie et al., 2013), our study revealed a subtle positive distractor effect in human choices. We further found that this weak positive distractor effect was indistinguishable between binary (distractor absent) and ternary trials (distractor present), indicating that distractor-based value distortions did not tap upon the immediate context. By interleaving a value estimation phase between the learning and choice phases, we found that subjective values were distorted already after the learning phase. These findings support the hypothesis that the distractor influence unfolds over a long timescale.

Context effects have been widely reported in psychology and behavioral economics in various domains including multiattribute consumer choices (Dumbalska et al., 2020; Webb et al., 2021) and risky choices (Chau et al., 2014, 2020; Mohr et al., 2017; Soltani et al., 2012). Typically, these violations are obtained in single-shot decisions, where participants need to compute the value of novel alternatives (based on subjective or externally determined criteria) on a single trial basis. Distractor effects, which is a certain type of context effects, have been reported both in single-shot decisions (Bavard & Palminteri, 2023; Cao & Tsetsos, 2023; Gluth et al., 2018, 2020; Itthipuripat et al., 2015; Louie et al., 2013) and in decisions that tap upon learned values. Finding that in our learning task the immediate context does not exert any influence on choices does not automatically overrule previous distractor effects obtained in single-shot tasks. It rather suggests that in value-learning tasks, where choices are guided by past rewards, the relative influence of the immediate context dissipates. This null immediate context effect could mean that the entire set of alternatives could be kept in memory and guide value computations, even in binary trials. Value-learning tasks that tax the memory by involving a larger number of alternatives could perhaps reveal a more complex interplay between the influence of the temporal and the immediate context.

Although the design of our study closely followed the design in Louie et al. (Louie et al., 2013), we did not replicate the strong negative distractor effect they found in the choice behavior of two monkeys. The negative distractor effect in Louie et al. was described as an immediate context effect attributed to divisive normalization. However, recent research (Cao & Tsetsos, 2023) has shown that a negative distractor effect can emerge without the recruitment of normalization mechanisms. This can occur when learning is conducted through choices among two or more alternatives, with reward feedback delivered exclusively for the chosen alternative. In these cases, in a high-distractor context, the distractor alternative is chosen (sampled) more frequently compared to a low-distractor context. As a consequence, participants in the high-distractor context sample the two high-value alternatives less often, exhibiting a lower choice accuracy due to increased value uncertainty (i.e., an “emergent” negative distractor effect). While in Louie et al. single-item learning trials preceded the choice phase, it is conceivable that the two monkeys continued to learn the values of the alternatives from the partial feedback received in the choice phase. This could have led to an “emergent” negative distractor effect owing to compromised learning in the high-distractor context (Cao & Tsetsos, 2023). The rapidly stabilized learning in our task prevented an “emergent” effect from arising. Thus, the discrepancy between our findings and those in Louie et al. (Louie et al., 2013) can be reconciled if the negative distractor effect was “emergent” in their case.

A curious finding in our study was that value estimates and choices disclosed seemingly different effects. In estimation, the distractor influence on value representations did not align with either classic positive or negative distractor effects. Instead, we found that the distractor value exerted a global subtractive influence on the estimated values of the two high-value alternatives (HV and LV). This subtractive effect in estimates together with the weak positive distractor effect in the choice task and could indicate separate mechanisms operating in the two tasks. However, in targeted analyses, we reported that value representations in the estimation and choice tasks were common. Consequently, we proposed a common value distortion mechanism in both choice and estimation. The discrepancy in the distractor influence in the two tasks could be reconciled if the estimation task is characterized by a larger response noise, effectively masking the overall small positive distractor effect. We formalized this mechanism using a stochastic rank-based model, inspired by decision-by-sampling theory (Noguchi & Stewart, 2018; Stewart et al., 2006). Although we favored this mechanism due to its parsimony, we acknowledge that less parsimonious schemes are conceivable. For example, both divisive and range normalization may at play, but the influence of each normalization scheme could vary across the estimation and choice phases. Future work could delve further into the potential interplay between these two prominent normalization mechanisms, also comparing hybrid schemes against more parsimonious ones.

To conclude, our findings suggest that during value-learning, the value of an inferior (distractor) alternative distorts value representations over a long timescale spanning several learning and choice trials. Crucially, the functional form of this distortion is incompatible with divisive and range normalization theories. Instead, it is best captured by a mechanism that constructs subjective value representations via comparing the value of the target alternative with the value of other alternatives, sampled from the memory context. These findings shed new light on the mechanisms that govern value learning and decision-making among multiple alternatives.

## Acknowledgments

This work was supported by the EU Horizon 2020 Research and Innovation Program (ERC starting grant no. 802905) to Konstantinos Tsetsos. The funders had no role in the study design, data collection and analysis, decision to publish, or preparation of the manuscript. We thank Yinan Cao for generously providing us with his insights during the early stage of this research.

## Author contributions

Maryam Tohidi-Moghaddam: Conceptualization, Methodology, Software, Validation, Formal Analysis, Investigation, Data Curation, Writing – Original Draft, Writing – Review & Editing, Visualization.

Konstantinos Tsetsos: Conceptualization, Methodology, Validation, Writing – Original Draft, Writing – Review & Editing, Supervision, Project Administration, Funding Acquisition.

## Supplemental Material

### Methods

#### Participants and Procedure

We collected data from thirty participants (22 female and 8 male, age range: 20-35). All participants had normal or corrected-to-normal vision were right-handed, with the exception of one participant who was left-handed. The experimental session lasted between 3 and 3.5 hrs and was broken down into 48 mini-blocks with each mini-block lasting ∼4 minutes. Participants had the opportunity to take short breaks (< 1 minute) between mini-blocks and one or two longer breaks (5-10 minutes) upon request. Participants received 30-35 EUR for their participation (hourly net rate = 10 EUR). In addition, we included a 10 EUR completion bonus and a maximum of 15 EUR task performance bonus to further incentivize participants to accurately perform the task. The average final task performance bonus was 13.76 EUR (95% CI = [13.05 EUR, 14.48 EUR], which was significantly higher than what would have been expected if they chose randomly (*t*_(29)_ = 41.3, *p* < .001).

#### Main task

The task was implemented using Psychophysics-3 Toolbox in Matlab (MathWorks, 2019) on a 21” monitor (1920 x 1080 px screen resolution and 120-Hz refresh rate). Stimuli were presented on a grey background, [127.5 127.5 127.5]. Participants viewed the monitor in a dimly lit room from a 60 cm distance. Participants completed 48 mini-blocks of a color-reward association task. They were instructed to imagine collecting colored coins to maximize their overall reward throughout the task. In each mini-block, participants initially learned the reward-value associations during the learning phase and later faced binary and ternary choices in the choice phase. The total reward obtained during the task was referred to as the task performance bonus.

Each mini-block was composed of 4 phases: 1-learning, 2-estimation (first/ pre-choice), 3-choice, and 4-estimation (second/post-choice) phases (Figure 2). The second and fourth phases followed identical procedures. In the learning phase, participants first serially learned the values associated with three colored alternatives. In the first estimation phase, they were asked to report the estimated value (i.e., subjective value) for each of those three learned alternatives by moving a slider on a bar. In the choice phase, they made choices among ternary and binary combinations of those alternatives. After the choice phase participants reported estimated values in a second and final estimation phase.

In each mini-block, there were three colored coins (red, blue, and yellow) randomly assigned to one of three values (HV, LV, DV, specific to each mini-block). The mapping between the colors and the values was fully counterbalanced (across six possible color-reward permutations). The combination of HV, LV, and DV alternative values represented the context that remained unchanged throughout each mini-block. We sampled the three values from the following sets HV = {55, 50}, LV = HV - 5, DV = {18,40}, which resulted in four unique value contexts. Combining these contexts with a full color-reward permutation, led to 24 unique possible mini-blocks (six color-reward permutations times four unique value contexts). These mini-blocks were repeated twice, in a total of 48 mini-blocks. In half of the mini-blocks, no reward feedback was provided during the choice phase (“No-Feedback” mini-blocks), while in the remaining half, the reward associated with each choice was displayed (“Feedback” mini-blocks). Participants were verbally informed that “Feedback” and “No-Feedback” mini-blocks were randomly interleaved, and they could not predict the type of mini-block before the choice phase. This design ensured that participants equally attended the learning phase in both “Feedback” and “No-Feedback” conditions. All participants received verbal instructions about the task structure and commenced the main experiment after completing two practice mini-blocks. Finally, all participants received verbal instructions on the structure of the task and started the main experiment after completing two practice mini-blocks.

#### Stimuli

##### Alternatives

The stimuli were three colored filled circles (diameter size = 1.5 visual degrees). The colors were yellow = [255 250 100], red = [255 90 90], and blue = [100 153 255].

##### Reward

The presented reward was a set of moving dots (diameter dot = .15 visual degrees) framed in a circle (frame thickness= .075 visual degrees, frame color = [51 51 51]). The number of dots was equal to the associated value of the chosen alternative (see the value sets mentioned in the main task section). The color of each dot was grayscale ([127.5 127.5 127.5]) and its contrast was drawn from a uniform distribution: [.6 1]. The reward presentation duration in both learning and “Feedback” choice trials was .75 seconds. We presented the dots in 10 consecutive frames (frame duration = 75 (ms)) and induced motion by randomly assigning each dot to a new location in each frame. The motion aimed to introduce uncertainty in the perceived numbers of dots, minimizing the possibility of participants learning the rewards after just one trial.

#### Mini-block structure

##### Learning phase

Participants completed 9 randomly presented learning trials (three per each alternative). Each trial started with the presentation of a fixation cue for a random short time (randomly drawn from a truncated exponential distribution: [.5 1] and mean = .75 (seconds)). Then the stimulus, one colored coin, was presented at the center of the screen for a maximum of 2 seconds. Participants were asked to click on the stimulus within 2 seconds to collect its associated reward. If they failed to do so, they missed the reward and saw a “Missed response” message on the screen. Once the stimulus appeared on the screen the mouse pointer appeared on a random position of an imaginary circle whose center aligned with the stimulus (diameter = 2 visual degrees). Immediately after clicking on the stimulus, participants could see a centrally presented cloud of moving representing the reward associated with the clicked stimulus. The reward was presented for .75 seconds and then followed by a gray screen with the fixation point on the center for 1.25 seconds as inter-trial-interval (ITI).

##### Estimation phase

We had 3 estimation trials, one for each alternative, presented in random order. In each estimation trial, we presented a target alternative (shown at the bottom-center of the bar) and asked participants to report the estimated value of the alternative corresponded to. They did so by first moving a slider on a bar within 10 seconds to find the best estimation and then pressing the “SPACE” bar to confirm their estimation. After 10 seconds, the current trial was automatically terminated and the next estimation trial was initiated. At the top of the two ends of the bar, we statically displayed the minimum (15 dots) and maximum (85 dots) possible number of dots, which we encouraged participants to use as reference. As participants adjusted the slider, they could instantly observe the number of moving dots within a set displayed at the top-center, changing in real-time.

##### Choice Phase

The choice phase included 12 ternary and 12 binary choice trials which were presented in random order. We had 3 combinations of binary trials {(HV, LV), (HV, DV), (LV, DV)} and presented each 4 times. The mapping between the location of the alternatives, (left, top, right) in ternary trials and (left, right) in binary trials, and the alternatives was counterbalanced. As in the learning trials, each trial began with the presentation of a fixation cue for a short delay (randomly drawn from a truncated exponential distribution: [.5 1] and mean = .75 (seconds)). Then the stimuli appeared on the screen for a maximum of 2 seconds. The stimuli were equidistant from the center (eccentricity = 2 visual degrees). Participants were asked to click on their preferred stimulus within 2 seconds to collect the associated reward for their choice. Failure to respond within the deadline resulted in a “Missed response” message appearing on the screen. On each choice trial the mouse pointer was initiated at the center of the screen. In “Feedback” mini-blocks, the reward was presented at the center of the screen immediately after a choice was made. The reward stayed on the screen for .75 seconds, followed by 1.25 seconds ITI. In “No-Feedback” mini-blocks a choice was directly followed by the 1.25 seconds ITI window.

##### Bonus window

We presented a bonus window twice, at the end of each estimation phase. This window illustrated the cumulative bonus earned from the start of the task until that moment, represented by a pink bar whose length visually reflected the accumulated bonus. A cyan line indicated the maximum possible bonus attainable. Consequently, the space between the end of the pink bar and the cyan represented the bonus participants had missed.

#### Behavioral Analyses

##### Choice prediction using subjective values

To convert the estimated values into pseudo-choices, we applied an *argmax* selection rule (like the one used here (Bavard & Palminteri, 2023)) on each ternary trial as:

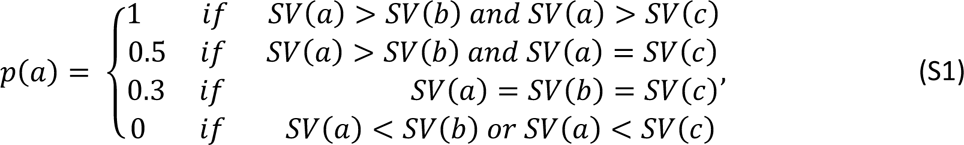

and on each binary trial as:

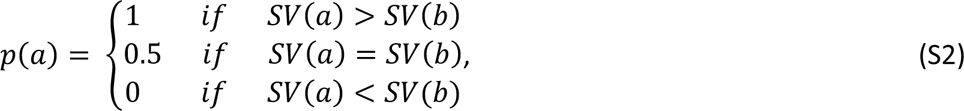

where *SV(x)* is the value estimate for alternative *x* and *p(x)* is the predicted choice probability of that alternative in each trial.

#### Computational Models

To model the choice behavior objective values (*OV_i_*) were transformed into “subjective” values (*SV_i_*), with the transformation type differing across models. Transformed values (*SV_i_*) were converted to choice probabilities by applying a probit function (using a numerical approach, Equation S3). For each alternative *i* the choice probability *p(i)* was calculated as:

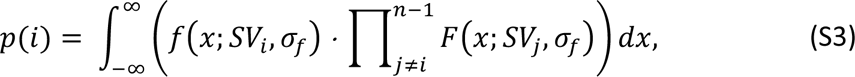

where *f* indicates the probability density function and *F* is the cumulative distribution function. Here we assumed that all represented subjective values are normally distributed (*N(SV,α^2^_f_))* and *n* is the number of alternatives. We used this range: [eps 100] for *α^2^_f_* to fit the model.

We used four computational models with each describing an alternative mechanism mapping objective values into their subjective (transformed) counterparts. ***Probit model (Pr)***: This model assumes that the subjective values should be unaffected by the choice-set context. Thus, there is no transformation in this model and the subjective values are encoded as equal to the objective values.

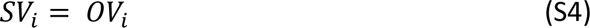

##### Divisive normalization model (DN)

The subjective value of each alternative is normalized by the sum of all alternatives.

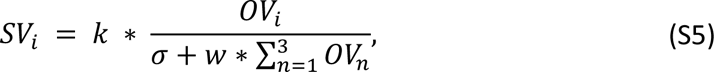

where *k*, *α*, and *w* are gain, semi-saturation, and weight terms. To fit the model we constrained parameters in these ranges, respectively: [1 200], [1 200], and [0 1].

##### Range normalization model (RN)

The subjective value of each alternative is normalized by the range of values in a given context.

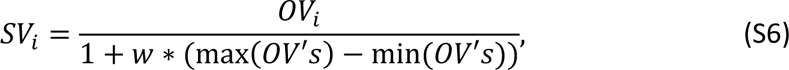

where *OV’s* refers to a vector consisting of all alternative values in the context and *w* is a weight term. To fit the model we constrained *w* between [0 1].

##### Rank-based model (RB)

This model implements relative value coding using a series of memory-based binary comparisons across all possible pairs of alternatives in the context. The subjective value of each alternative is the total number of times that this alternative was the winner across all memory-based binary comparisons.

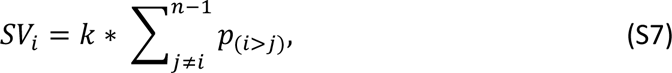

where *k* represents the total number of binary comparisons, constrained in the [1 200] range. *p_(i>j)_* is the probability that alternative *i* is deemed as better than alternative *j* in a given binary comparison. We derived this probability using a softmax function:

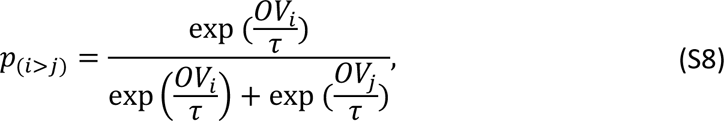

where *ι−* is the temperature parameter, which was constrained in the [1 100] range when fitting the model.

##### Model simulations

The model predictions shown in Figure 1 were generated using numerical approximations (see Equation S3). In all models, we used the same value sets: HV = 50, LV = 45, and DV = {0, 7, 13, 20, 27, 33, 40}. The parameters for each model were fixed as follows: Probit: *α^2^_f_* = 30.25, divisive normalization: *α^2^_f_* = 9, k = 100, *σ* = 50 and *w* = 1, and range normalization: *α^2^_f_* = .25, and *w* = 1.

##### Model fitting procedure

We fitted the models by minimizing the negative log-likelihood (*NLL*) summed over all choice trials (including all ternary and binary trials) (Wilson & Collins, 2019). For each participant, we fitted each model once on the entire set of choice trials, including both “Feedback” and “No-Feedback” mini-blocks (∼1152 choice trials). We optimized the parameters using Matlab’s *fmincon.m* function (MaxFunEvals = 5000; MaxIter = 5000; TolFun = 1e-20; TolX = 1e-20). To avoid local optima, we refitted each model for each participant 10 times using a grid of randomly generated starting values for the free parameters.

##### Model selection

We used a fixed-effects and a random-effects model comparison. We used the Bayesian information criterion (*BIC*) to compare different models. The *BIC* is quantified as below:

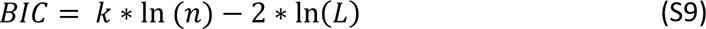

where *L* is the likelihood value obtained from the model fit to all choice data, *k* is the number of free parameters in each model and *n* is the number of trials we used to fit the model (Schwarz, 1978). We computed the *BIC* score for each participant and each model. Then for each participant the model with the lowest *BIC* was marked as the best-individualized model. Finally, we reported the *ΔBIC* of each model relative to the best-individualized model (Figure 6B).

We also fitted the models using a 6-fold cross-validation procedure. To do so, for each participant, we split the trials into 6 parts (i.e., folds) and then fitted each model to a “training” set (comprising five random folds). We used the best-fitted parameters of the “training” set to calculate the *LL* summed across trials in the left-out “test” fold. We repeated this process over test folds (6 times) and the final cross-validated *LL* was computed as the mean *LL* across cross-validated folds. We then used this mean *LL* to calculate each model’s posterior frequency and protected exceedance probability (i.e., the probability corrected for the chance level that a model is more likely than any others) using the variational Bayesian analysis (VBA) toolbox (Daunizeau et al., 2014; Rigoux et al., 2014).

## Results

**Figure S1.**
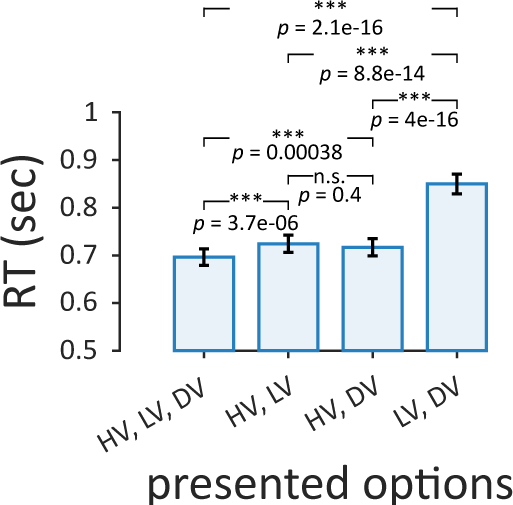
Reaction time of choice phase. The reaction time of participants’ choices (both correct and incorrect choices) illustrated for each trial type. The RT in ternary trials is significantly lower than any binary trials. Within binary trial types, “LV, DV” pair has the significant higher RT than the other two binary pairs, the error bars are standard errors of the mean across participants.

**Figure S2.**
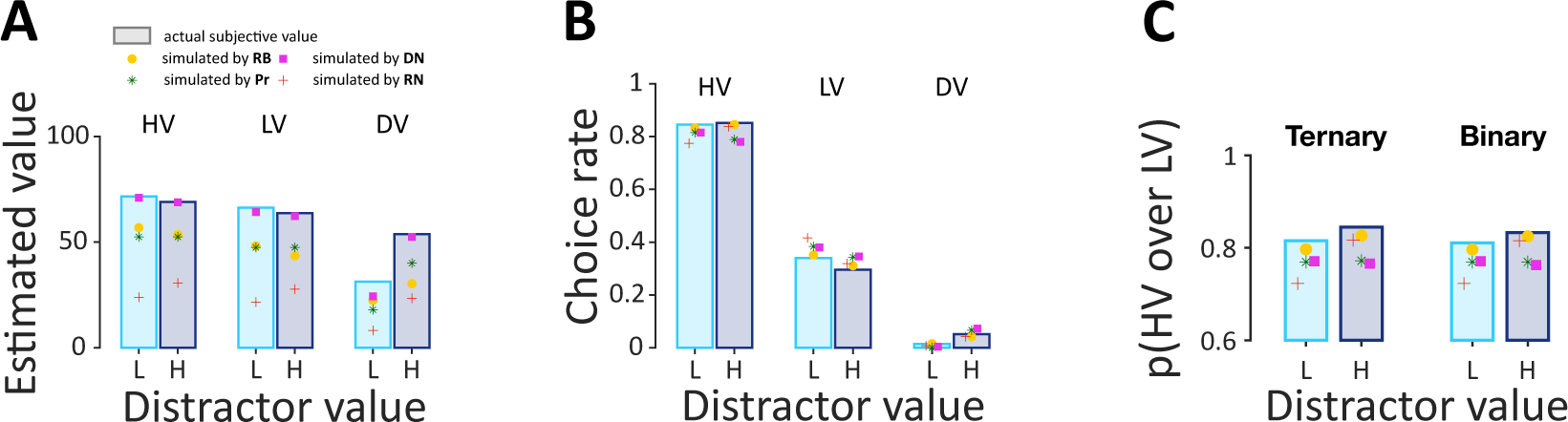
Model predictions. **(A)** The simulated estimated value of each alternative of each model superimposed on the actual estimated value from the first estimation phase. **(B)** The simulated choice rate of each model superimposed on the actual choice. **(F)** The simulated relative probability of HV over LV of each model superimposed on the actual relative probability. It shows that RB and RN predicted the positive distractor effect, DN predicted the negative distractor effect and Pr predicted the null distractor effect. In all panels, the simulated data points were calculated using the best-fitted parameters of each model, and the error bars are standard errors of the mean across participants.

**Table S1.** Best-fitted model parameters, (mean ± standard error of the mean across participants).

| **Model** | RB | Pr | DN | RN |
| --- | --- | --- | --- | --- |
| **Parameter** |  |  |  |  |
| **k** | 42.46±9.36 | - | 122.63±8.57 | - |
| **$\tau$** | 21.39±6.57 | - | - | - |
| **$\sigma$** | - | - | 78.57±10.72 | - |
| **w** | - | - | 0.21±0.07 | 0.22±0.06 |
| **$\sigma^2_f$** | 53.67±3.44 | 33.47±4.96 | 48.18±5.27 | 9.13±1.92 |

## References

Bavard, S., & Palminteri, S. (2023). The functional form of value normalization in human reinforcement learning. eLife, 12, e83891. 10.7554/eLife.83891

Berkowitsch, N. A. J., Scheibehenne, B., & Rieskamp, J. (2014). Rigorously testing multialternative decision field theory against random utility models. Journal of Experimental Psychology. General, 143(3), 1331–1348. 10.1037/a0035159

Cao, Y., & Tsetsos, K. (2022). Clarifying the role of an unavailable distractor in human multiattribute choice. eLife, 11, e83316. 10.7554/eLife.83316

Cao, Y., & Tsetsos, K. (2023). *Decision bias and sampling asymmetry in reward-guided learning* [Preprint]. Neuroscience. 10.1101/2023.09.10.557023

Chang, L. W., Gershman, S. J., & Cikara, M. (2019). Comparing value coding models of context-dependence in social choice. Journal of Experimental Social Psychology, 85, 103847. 10.1016/j.jesp.2019.103847

Chau, B. K., Kolling, N., Hunt, L. T., Walton, M. E., & Rushworth, M. F. S. (2014). A neural mechanism underlying failure of optimal choice with multiple alternatives. Nature Neuroscience, 17(3), Article 3. 10.1038/nn.3649

Chau, B. K., Law, C.-K., Lopez-Persem, A., Klein-Flügge, M. C., & Rushworth, M. F. (2020). Consistent patterns of distractor effects during decision making. eLife, 9, e53850. 10.7554/eLife.53850

Daunizeau, J., Adam, V., & Rigoux, L. (2014). VBA: A Probabilistic Treatment of Nonlinear Models for Neurobiological and Behavioural Data. PLoS Computational Biology, 10(1), e1003441. 10.1371/journal.pcbi.1003441

Dumbalska, T., Li, V., Tsetsos, K., & Summerfield, C. (2020). A map of decoy influence in human multialternative choice. Proceedings of the National Academy of Sciences of the United States of America, 117(40), 25169–25178. 10.1073/pnas.2005058117

Frydman, C., & Jin, L. J. (n.d). Efficient Coding and Risky Choice.

Gluth, S., Kern, N., Kortmann, M., & Vitali, C. L. (2020). Value-based attention but not divisive normalization influences decisions with multiple alternatives. Nature Human Behaviour, 4(6), Article 6. 10.1038/s41562-020-0822-0

Gluth, S., Spektor, M. S., & Rieskamp, J. (2018). Value-based attentional capture affects multi-alternative decision making. eLife, 7, e39659. 10.7554/eLife.39659

Greene, W. H. (2000). Econometric Analysis (4th edition). Prentice Hall.

Hayes, W. M., & Wedell, D. H. (2023). Effects of blocked versus interleaved training on relative value learning. Psychonomic Bulletin & Review. 10.3758/s13423-023-02290-6

Holper, L., Brussel, L. D. V., Schmidt, L., Schulthess, S., Burke, C. J., Louie, K., Seifritz, E., & Tobler, P. N. (2017). Adaptive Value Normalization in the Prefrontal Cortex Is Reduced by Memory Load. eNeuro, 4(2). 10.1523/ENEURO.0365-17.2017

Itthipuripat, S., Cha, K., Rangsipat, N., & Serences, J. T. (2015). Value-based attentional capture influences context-dependent decision-making. Journal of Neurophysiology, 114(1), 560–569. 10.1152/jn.00343.2015

JASP Team. (2023). *JASP (Version 0.18.0)[Computer software]* (0.18.0) [Computer software]. https://jasp-stats.org/

Juechems, K., Altun, T., Hira, R., & Jarvstad, A. (2022). Human value learning and representation reflect rational adaptation to task demands. Nature Human Behaviour, 6(9), Article 9. 10.1038/s41562-022-01360-4

Kass, R. E., & Raftery, A. E. (1995). Bayes Factors. Journal of the American Statistical Association, 90(430), 773–795. 10.2307/2291091

Khaw, M. W., Glimcher, P. W., & Louie, K. (2017). Normalized value coding explains dynamic adaptation in the human valuation process. Proceedings of the National Academy of Sciences, 114(48), 12696–12701. 10.1073/pnas.1715293114

Kohl, C., Wong, M. X., Wong, J. J., Rushworth, M. F., & Chau, B. K. (2023). Intraparietal stimulation disrupts negative distractor effects in human multi-alternative decision-making. eLife, 12, e75007. 10.7554/eLife.75007

Kontek, K., & Lewandowski, M. (2018). Range-Dependent Utility. Management Science, 64(6), 2812–2832. 10.1287/mnsc.2017.2744

Louie, K., & De Martino, B. (2014). Chapter 24—The Neurobiology of Context-Dependent Valuation and Choice. In P. W. Glimcher & E. Fehr (Eds.), Neuroeconomics (Second Edition) (pp. 455–476). Academic Press. 10.1016/B978-0-12-416008-8.00024-3

Louie, K., & Glimcher, P. W. (2012). Efficient coding and the neural representation of value. Annals of the New York Academy of Sciences, 1251(1), 13–32. 10.1111/j.1749-6632.2012.06496.x

Louie, K., Khaw, M. W., & Glimcher, P. W. (2013). Normalization is a general neural mechanism for context-dependent decision making. Proceedings of the National Academy of Sciences, 110(15), 6139–6144. 10.1073/pnas.1217854110

Luce, R. D. (1959). Individual choice behavior (pp. xii, 153). John Wiley.

Luce, R. D. (1977). The choice axiom after twenty years. Journal of Mathematical Psychology, 15(3), 215–233. 10.1016/0022-2496(77)90032-3

Madan, C. R., Spetch, M. L., Machado, F. M. D. S., Mason, A., & Ludvig, E. A. (2021). Encoding Context Determines Risky Choice. Psychological Science, 32(5), 743– 754. 10.1177/0956797620977516

Mohr, P. N. C., Heekeren, H. R., & Rieskamp, J. (2017). Attraction Effect in Risky Choice Can Be Explained by Subjective Distance Between Choice Alternatives. Scientific Reports, 7(1), Article 1. 10.1038/s41598-017-06968-5

Noguchi, T., & Stewart, N. (2014). In the attraction, compromise, and similarity effects, alternatives are repeatedly compared in pairs on single dimensions. Cognition, 132(1), 44–56. 10.1016/j.cognition.2014.03.006

Noguchi, T., & Stewart, N. (2018). Multialternative decision by sampling: A model of decision making constrained by process data. Psychological Review, 125(4), 512–544. 10.1037/rev0000102

Paetz, F., & Steiner, W. J. (2018). Utility independence versus IIA property in independent probit models. Journal of Choice Modelling, 26, 41–47. 10.1016/j.jocm.2017.06.001

Palminteri, S., Khamassi, M., Joffily, M., & Coricelli, G. (2015). Contextual modulation of value signals in reward and punishment learning. Nature Communications, 6(1), Article 1. 10.1038/ncomms9096

Palminteri, S., & Lebreton, M. (2021). Context-dependent outcome encoding in human reinforcement learning. Current Opinion in Behavioral Sciences, 41, 144–151. 10.1016/j.cobeha.2021.06.006

Rangel, A., & Clithero, J. A. (2012). Value normalization in decision making: Theory and evidence. Current Opinion in Neurobiology, 22(6), 970–981. 10.1016/j.conb.2012.07.011

Rigoli, F. (2019). Reference effects on decision-making elicited by previous rewards. Cognition, 192, 104034. 10.1016/j.cognition.2019.104034

Rigoux, L., Stephan, K. E., Friston, K. J., & Daunizeau, J. (2014). Bayesian model selection for group studies—Revisited. NeuroImage, 84, 971–985. 10.1016/j.neuroimage.2013.08.065

Soltani, A., Martino, B. D., & Camerer, C. (2012). A Range-Normalization Model of Context-Dependent Choice: A New Model and Evidence. PLOS Computational Biology, 8(7), e1002607. 10.1371/journal.pcbi.1002607

Stewart, N., Chater, N., & Brown, G. D. A. (2006). Decision by sampling. Cognitive Psychology, 53(1), 1–26. 10.1016/j.cogpsych.2005.10.003

The MathWorks Inc. (2022). MATLAB version: 9.12.0 (R2022a) [Computer software]. https://www.mathworks.com

Webb, R., Glimcher, P. W., & Louie, K. (2020). Divisive normalization does influence decisions with multiple alternatives. Nature Human Behaviour, 4(11), Article 11. 10.1038/s41562-020-00941-5

Webb, R., Glimcher, P. W., & Louie, K. (2021). The Normalization of Consumer Valuations: Context-Dependent Preferences from Neurobiological Constraints. Management Science, 67(1), 93–125. 10.1287/mnsc.2019.3536

## References

Schwarz, G. (1978). Estimating the Dimension of a Model. The Annals of Statistics, 6(2), 461–464.

Wilson, R. C., & Collins, A. G. (2019). Ten simple rules for the computational modeling of behavioral data. eLife, 8, e49547. 10.7554/eLife.49547

